# Human iPSC-derived retinal pigment epithelium: a model system for identifying and functionally characterizing causal variants at AMD risk loci

**DOI:** 10.1101/440230

**Authors:** Erin N. Smith, Agnieszka D’Antonio-Chronowska, William W. Greenwald, Victor Borja, Lana R. Aguiar, Robert Pogue, Hiroko Matsui, Shyamanga Borooah, Matteo D’Antonio, Radha Ayyagari, Kelly A. Frazer

## Abstract

We evaluate whether human induced pluripotent stem cell-derived retinal pigment epithelium (iPSC-RPE) cells can be used to prioritize and functionally characterize causal variants at age-related macular degeneration (AMD) risk loci. We generated iPSC-RPE from six subjects and show that they have morphological and molecular characteristics similar to native RPE. We generated RNA-seq, ATAC-seq, and H3K27ac ChIP-seq data and observe high similarity in gene expression and enriched transcription factor motif profiles between iPSC-RPE and human fetal-RPE. We performed fine-mapping of AMD risk loci by integrating molecular data from the iPSC-RPE, adult retina, and adult RPE, which identified rs943080 as the probable causal variant at *VEGFA*. We show that rs943080 is associated with altered chromatin accessibility of a distal ATAC-seq peak, decreased overall gene expression of *VEGFA*, and allele specific expression of a non-coding transcript. These results provide insight into the mechanism underlying the association of the *VEGFA* locus with AMD.

## Introduction

AMD is a leading cause of vision loss that affects 1.6 million people over the age of 50 in the US (CDC) and has limited therapeutic options (Al-Zamil and Yassin, 2017). Disease development manifests in progressive degeneration in response to oxidative stress and inflammation of the retinal pigment epithelium (RPE) (Kinnunen et al., 2012), a monolayer consisting of a few million cells most densely located in the macula of the eye (Panda-Jonas et al., 1996). AMD has a strong genetic component (Seddon et al., 2005), and through a large international study of 16,144 AMD cases and 17,832 controls, 52 independent AMD risk-variants mapping to 34 AMD-associated loci have been identified (Fritsche et al., 2016). As in many common diseases (Gusev et al., 2014; Maurano et al., 2012), the majority of these loci have strongly associated variants in noncoding regions of the genome, suggesting that they may act through gene regulation. However, while candidate target genes have been identified (Fritsche et al., 2016), the causal variants and their gene targets are generally unknown.

Regulatory genetic variation is often cell-type specific and can be studied through genetic analysis of molecular traits such as gene regulation and expression (Albert and Kruglyak, 2015). However, characterizing genetic variation in human RPE is challenging because the number of RPE cells in the human eye is limited (Panda-Jonas et al., 1996), can be affected by lifetime environmental exposures, and requires invasive procedures to collect samples. Induced pluripotent stem cell-derived RPE (iPSC-RPE) is a promising alternative to human RPE for genetic studies as a virtually unlimited number of cells can be obtained with the same genetic background non-invasively. iPSC-RPE has been shown to display characteristics of mature human RPE including polygonal and pigmented morphology, polarity of protein expression and secretion, phagocytosis of photoreceptor outer segments, and maintenance of RPE phenotypes after transplantation into mouse retina (Maruotti et al., 2015). Additionally, stem cell-derived RPE have been effectively transplanted into rodent and primate models, supporting their relevance *in vivo* (Davis et al., 2017; Kamao et al., 2014; Stanzel et al., 2014). Thus, iPSC-RPE could be an effective model system for functionally characterizing regulatory variation associated with AMD.

The identification and functional characterization of causal genetic variants has been improved through fine-mapping algorithms that can incorporate diverse epigenetic annotations. For example, accessible chromatin and active regulatory regions such as promoters and enhancers marked by histone 3 lysine-27 acetylation (H3K27ac) have been shown to be enriched for genetic variants associated with human diseases in cell types relevant for disease and can improve the prioritization of GWAS causal variants through fine-mapping strategies (Pickrell, 2014). Additionally, while many GWAS loci harbor genes that have been implicated in AMD, such as *VEGFA* which encodes the VEGF protein that is targeted by three current treatments for AMD (Gragoudas et al., 2004; Heier et al., 2012; Rosenfeld et al., 2006), the causal risk variant and the mechanism of increased disease risk is not known. Thus, the molecular characterization of gene expression and regulatory regions in iPSC-RPE could improve fine-mapping of AMD and lead to insights into mechanisms underlying genetic risk variants.

To investigate the utility of iPSC-RPE as a model system to characterize AMD risk variants, we generated iPSC-RPE from six human subjects and integrated gene expression, chromatin accessibility, and H3K27ac ChIP-seq data with complementary published data from human adult subjects to identify potential causal variants at AMD risk loci. We show that the iPSC-RPE shows morphological and molecular characteristics that are similar to native RPE including a characteristic polygonal shape, strong melanin pigmentation and expression, and strong ZO-1, BEST1, and MITF immunostaining. We show that iPSC-RPE gene expression profiles are highly similar to human fetal RPE, and that their ATAC-seq peaks are enriched for relevant transcription factor motifs. We performed fine-mapping of AMD risk loci integrating the molecular data from iPSC-RPE, human fetal RPE, and published human retina and RPE samples. At one locus, *VEGFA*, we show that the rs943080 risk allele is associated with regulatory protein binding in iPSC-RPE in a potentially disease-dependent manner, and that the risk allele results in decreased overall *VEGFA* expression, potentially through regulation by a non-coding transcript. These results establish a molecular hypothesis for the *VEGFA* genetic risk locus on AMD and illustrate the potential of iPSC-RPE as a model system to study the molecular function of genetic variation associated with AMD.

## Results

### Derivation of iPSC-RPE

To derive iPSC-RPE, we chose six iPSC lines from unrelated individuals of African American (1), European (3), and East Asian (2) ancestry in iPSCORE that have been established as being pluripotent with low levels of somatic variation (Panopoulos et al., 2017). We then applied a slightly modified version of a protocol validated by Maruotti et al. (Maruotti et al., 2015) (**Figure 1A**) to differentiate iPSC-RPE. The first pigmented foci of characteristic polygonal cells appeared after ~2-3 weeks, and virtually all cells were strongly pigmented by day 84 (**Figure 1B**). We assessed the purity of all six iPSC-RPE samples at day 84 by flow cytometry and observed that a high fraction of cells were ZO-1 and MITF double positive (mean = 90.8%, range = 88-98.1) (**Figure 1C**). We also examined the iPSC-RPE samples for characteristic RPE marker genes and observed membrane expression of ZO-1 and BEST1 and nuclear expression of MITF (**Figure 1D**). Thus, the iPSC-RPE displayed many of the morphological and molecular characteristics of RPE, confirming the robustness of the differentiation protocol.

**Figure 1:**
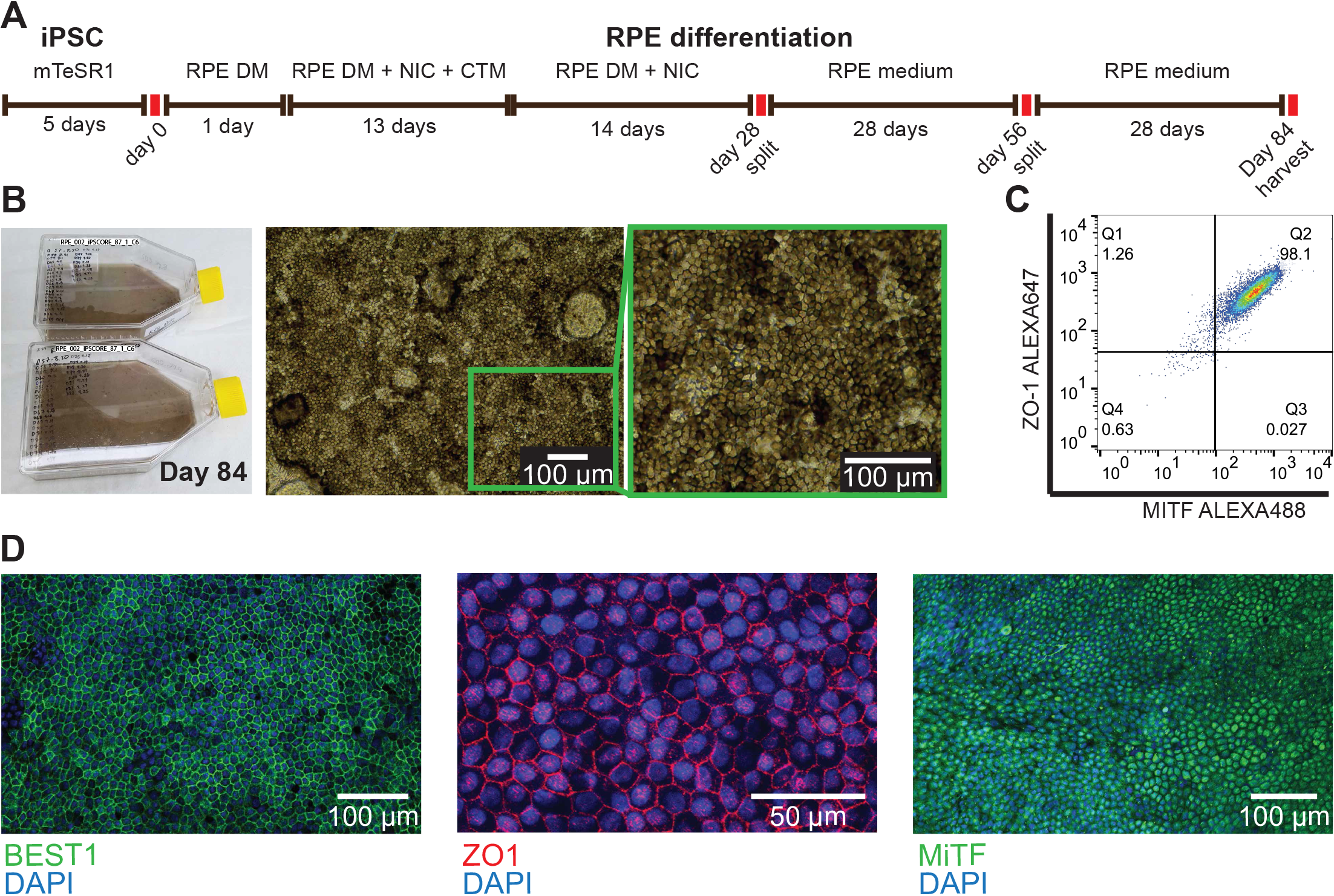
**A)** iPSC-RPE differentiation protocol. **B)** T150 flasks containing iPSC-RPE (iPSCORE 87_1) at day 84 (left). Bright-field image of iPSC-RPE sample (iPSCORE_42_1) at day 84 illustrating a highly-organized monolayer with strong melanin pigmentation and characteristic polygonal shape (right). **C)** Flow cytometry analysis of iPSC-RPE (iPSCORE_42_1) at day 84 showing high co-staining of Zonula Occludens 1 (ZO-1) and Microphthalmia-associated transcription factor (MITF). **D)** Immunofluorescence analysis of Bestrofin 1 (BEST1) (iPSCORE_29_1), ZO-1 (iPSCORE_29_1), and MITF (iPSCORE_42_1).

### The transcriptomes of human iPSC-RPE are similar to fetal RPE

We first compared iPSC-RPE gene expression to fetal RPE gene expression to examine their similarity overall and at previously established RPE signature genes. We generated RNA-seq data for the six iPSC-RPE, as well as one RPE obtained from a human fetal sample (**Table S1**). Using principal component analysis (PCA), we examined global trends between the iPSC-RPE and the fetal RPE RNA-seq in contrast to 222 iPSC lines (DeBoever et al., 2017) and 144 iPSC-derived cardiomyocytes (iPSC-CMs) (unpublished data) for which we have previously generated RNA-seq. The first two PCs split the iPSCs from the differentiated cells (iPSC-CMs and RPE), while PCs 3 and 4 were associated with strong clustering of the RPE samples (**Figure 2 A,B**). We therefore examined the top 1000 genes with the strongest weights associated with PC3 and 4 with gene ontology enrichment analysis to determine their associations with RPE-related functions. We observed enrichment for a number of RPE functions including sensory organ development (GO:0007423, P=5.2×10^−11^, PC3) and sensory perception of light stimulus (GO:0050953, P=2.0×10^−19^, PC4) (**Table S2**). We also found that genes associated with PC3 and PC4 were significantly enriched for previously identified RPE signature genes (Strunnikova et al., 2010) (P=4.6×10^−7^ and 3.8×10^−20^, respectively) and the expression patterns of these genes in iPSC-RPE were similar to the fetal RPE (**Figure 2C**). As a subset of these signature genes has been previously reported to be different between fetal RPE and stem-cell derived RPE (Liao et al., 2010), we examined their clustering pattern, but did not observe differential clustering (**Figure 2C**, “fetal, not stem-RPE”). It is possible that the high genomic integrity of the currently studied iPSC lines or improvements to the differentiation protocol (foci formation vs. a continuous monolayer) could have resulted in iPSC-RPE that were more similar to fetal RPE. Further, gene expression was highly correlated between fetal-RPE and iPSC-RPE across RPE signature genes (**Figure 2D**), suggesting that iPSC-RPE have transcriptomes that are highly similar to fetal RPE.

**Figure 2:**
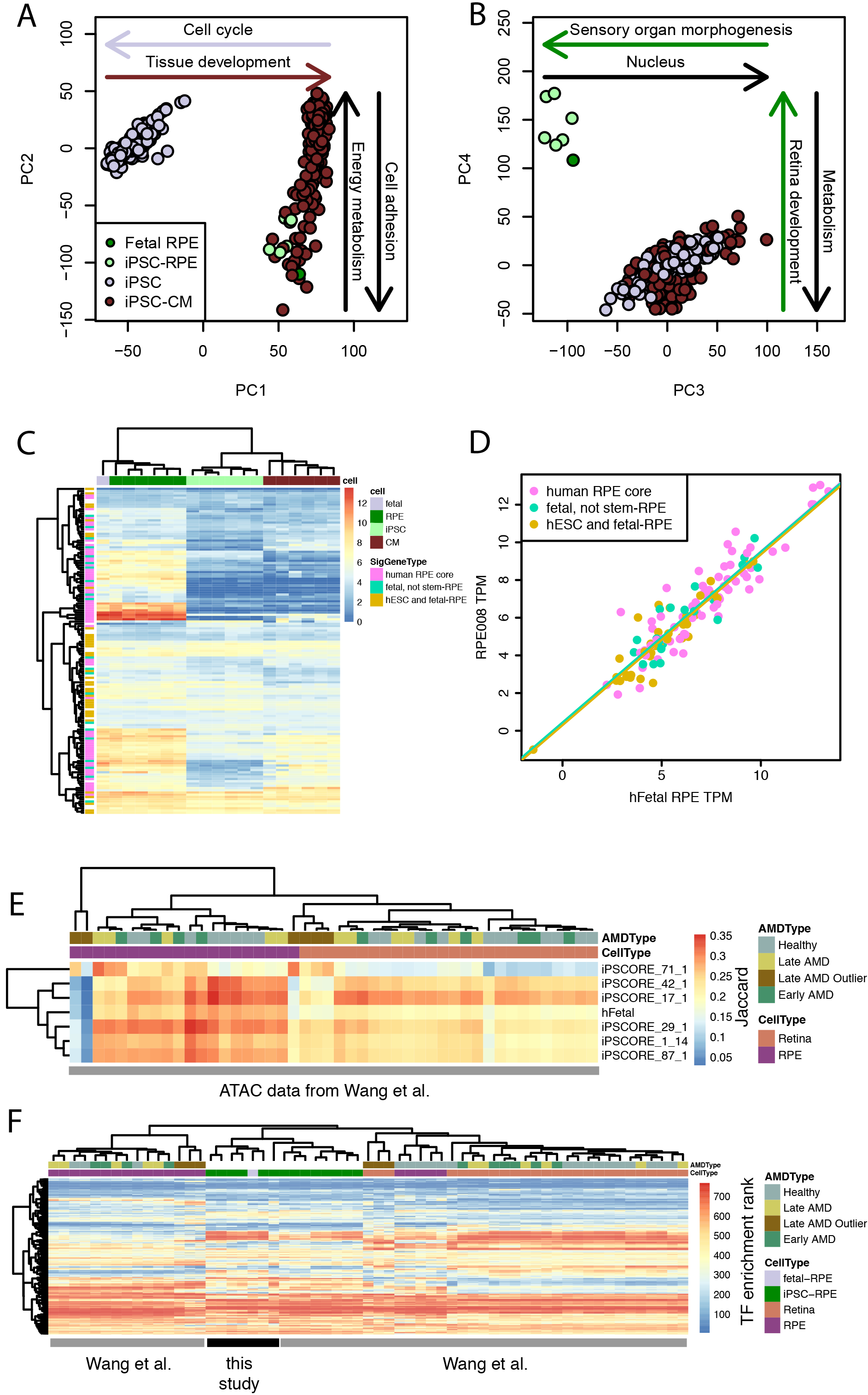
**A)** PCA plot of first two RNA-seq principal components (PCs) based on RNA-seq from 10,000 of the most variable genes. Annotated arrows indicate gene ontology enrichment for genes with in the top or bottom 10% weights for each PC. **B)** PCA of the third and fourth PCs showing the clustering of the iPSC-RPE samples with the fetal RPE sample. **C)** Heatmap showing expression levels of RPE signature genes (Strunnikova et al., 2010) in fetal RPE, iPSC-RPE, iPSC, and iPSC-derived cardiomyocytes. Genes are labeled by their gene group as annotated in (Liao et al., 2010). **D)** Scatterplot showing RNA-seq TPM for an iPSC-RPE (iPSCORE_1_14) compared to the fetal RPE sample for RPE signature genes. Points are color coded according to their classification in (Liao et al., 2010) and lines indicate linear regression best fits. **E)** Heatmap showing six iPSCORE iPSC-RPE (iPSCORE_42_1, 17_1, 29_1, 1_14 and 87_1) and human fetal RPE ATAC-seq peaks show stronger Jaccard similarity to native RPE ATAC-seq peaks than retina ATAC-seq peaks from adult eyes with and without AMD (Wang et al., 2018). Columns are color coded according to their AMD type and their cell type. Three RPE and three Retina samples from the same subject with late AMD were not similar to respectively the other RPE and Retina samples and were labeled as “Late AMD Outlier”. **F)** Heatmap showing similarity of transcription factor binding site enrichment in ATAC-seq peaks across samples. Color indicates the TF’s rank of enrichment within each sample with red indicating stronger enrichment. Columns are color coded according to their AMD type and their cell type.

### Chromatin accessibility of iPSC-RPE

We next examined whether the chromatin accessibility profiles of the iPSC-RPE were similar to fetal or adult RPE. We generated ATAC-seq data for the six iPSCORE iPSC-RPEs, and for the fetal RPE, (**Table S1**) and compared it to published RPE and whole retina tissues from adults with and without AMD (Wang et al., 2018). We compared the overlap of the accessible chromatin regions in iPSC-RPE or fetal RPE with adult RPE from both control and AMD subjects using the Jaccard similarity metric and observed high similarity (**Figure 2E**). We noted, however, that samples from the Wang et al. study derived from subject “Donor 1” appeared to be outliers with respect to the rest of the data. We therefore labeled these six samples as “Late AMD Outlier”. We then examined whether the accessible regions displayed similar transcription factor motif enrichments across all samples using HOMER (Heinz et al., 2010). We observed high enrichment for TF motifs important for RPE development and function in all RPE samples including OTX2, CRX, and SOX9 (**Table S3**). We compared the overall pattern of TF enrichment of the fetal and iPSC-RPE to those from Wang et al. by clustering the profiles by their enrichment rank in order to account for variation in binding specificity across studies. We observed that the iPSC-RPE from this study were most similar to the human fetal RPE and iPSC-RPE samples in Wang et al. (**Figure 2F**). Additionally, the fetal and stem-cell derived RPE showed stronger similarity to adult RPE samples than adult retina samples. These results suggest that iPSC-RPE show a fetal-like regulatory program that is similar to, but distinct from adult RPE, and different from adult retina.

### Enrichment of regulatory regions for AMD genetic risk

To establish whether regulatory regions in iPSC-RPE could be used to help identify potential causal variants for AMD, we generated H3K27ac ChIP-seq data from the iPSC-RPE and tested for enrichment of these and the chromatin accessibility regions for AMD genetic risk (Fritsche et al., 2016) using fgwas (Pickrell, 2014). We found that H3K27ac peak regions from fetal RPE, as well as ATAC-seq peak regions from iPSC-RPE, fetal RPE, adult AMD RPE, adult healthy RPE, and adult healthy retina, were enriched for AMD genetic risk (**Figure 3A**). In addition, exon regions and missense variants were also enriched. However, ATAC-seq regions from adult AMD (early and late) retina samples, H3K27ac peak regions from iPSC-RPE, and promoters showed positive associations, but were not significantly enriched. We also observed that the regions from the Late AMD Outlier samples showed depletion for AMD risk and we therefore excluded these samples from downstream analyses (**Figure S1**). Further, iPSC-RPE ATAC-seq and H3K27ac ChIP-seq regions from subjects of different ethnicities showed overall similar levels of enrichment and were therefore merged (**Figure S1**). We created a combined model of all annotations (**Figure 3A**), which removes redundant annotations and assesses the independent risk of each annotation, and observed that fetal-, iPSC-, and adult-RPE samples all provided independent positive information for AMD risk (**Figure 3B**). ATAC-seq regions from retina samples, however, showed negative enrichments, suggesting that the positive associations in the single model were captured by the other annotations and were RPE-specific. These results indicate that chromatin accessibility regions in RPE from adults, fetal, and iPSC-derived samples capture complementary risk regions associated with AMD risk and could be used to improve identification of potential causal variants.

**Figure 3:**
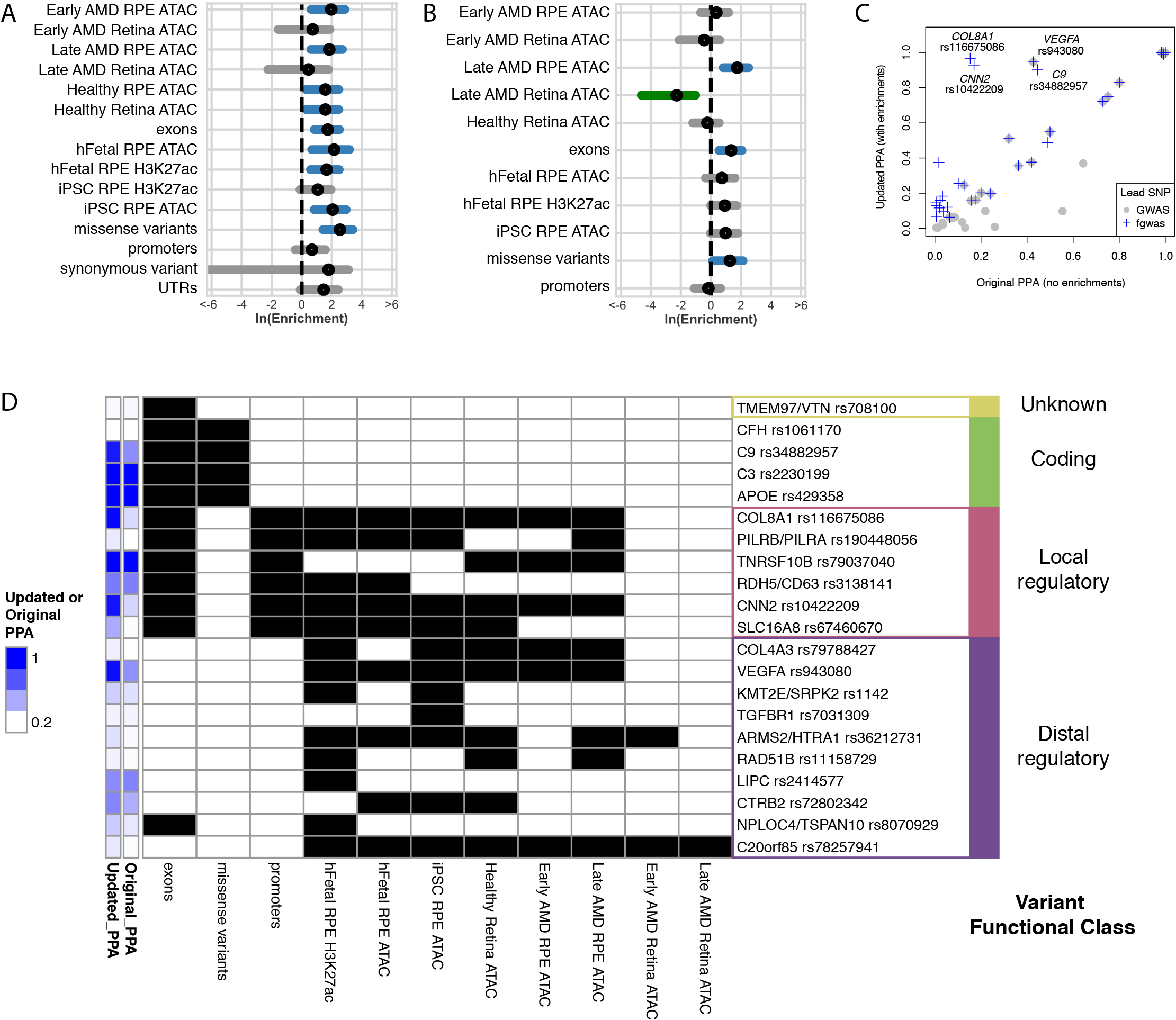
**A)** fgwas single enrichment of GWAS signal for multiple genome annotations. Black circles indicate ln(OR) and blue or grey lines show 95% CIs. Blue indicates a significant positive enrichment. **B)** Final combined model used by fgwas to prioritize variants. Black circles indicate ln(OR) and blue, grey, or green lines show 95% CIs (colored to indicate significance different from 0; green negative, blue positive). Note that in this combined model, associations reflect independent associations for each annotation and exclude annotations that did not significantly improve the model, and so differ from single enrichments shown in B. **C)** Scatterplot showing posterior probability of association (updated PPA) vs. original posterior probability of association (original PPA) for GWAS loci. GWAS lead SNPs are shown in grey. SNPs that were chosen as variant at the locus with the highest posterior probability of association by fgwas are shown in blue. When the variant with the highest updated PPA is the GWAS lead variant, the points overlap. The four variants where the fgwas prediction improved the PPA from < 0.5 to > 0.8 are labeled by their candidate gene and fgwas lead SNP. **D)** Heatmap indicating the presence (black) or absence (white) of each annotation for the 21 lead fgwas variants that were prioritized by the model. The original and updated PPAs are shown in blue and are the same as in C. Variants are grouped into variant functional classes according to whether they were unknown, coding, local regulatory (located in promoters), or contained in distal regulatory (ATAC-seq or H3K27ac w/o promoter) annotations.

### Fine-mapping AMD-associated loci with fgwas

To identify potentially causal variants associated with AMD, we used fgwas to perform fine-mapping of 32 of the 34 AMD risk loci for which we were able to obtain sufficient data (see Experimental Procedures) based on the estimates from the combined model in Figure 3B. For nine of the 32 risk loci, we identified a single variant with a posterior probability of association (PPA) > 0.8 (**Table S4**). Of these, four showed strong shifts between their PPAs before and after updating with the enrichments, including three loci where the GWAS lead variant was not the prioritized variant (*COL8A1*, *CNN2*, and *C9*), and one where the lead variant was the prioritized variant (*VEGFA*) (**Figure 3C**). To obtain insight into the types of regulation that could be associated with each GWAS locus, we examined how each of the top variants prioritized by fgwas was associated with each annotation and classified each into putative functional classes according to their annotation patterns (**Figure 3D**). Of the 32 fgwas lead variants, 21 were associated with an annotation. The majority of the variants (16/21 (76%)) were located in regulatory regions, overlapping both promoter (local regulatory, 6/21 (29%)) and proximal regions (distal regulatory, 10/21 (48%)). An additional four were missense variants (coding, 4/21 (19%)). One variant was located in an exon (in a 3’UTR), but was not associated with other annotations, and was classified as unknown (1/21, (5%)). We observed 13 regulatory variants with ATAC-seq annotations present in either iPSC-RPE or adult AMD RPE (early or late-stage AMD), of which 7/13 were present in both sample types. The remainder were present only in iPSC-RPE (4/13) or only in adult AMD RPE (2/13), suggesting that the majority of ATAC-seq regions associated with AMD GWAS loci are present in iPSC-RPE, but that in some cases they are only present in RPE from subjects with AMD. Overall, these results establish a set of prioritized variants at AMD loci, identify four variants with high PPA that were not previously distinguished from other variants at their loci, and illustrate that iPSC-RPE and adult RPE provide overlapping but complementary regulatory annotations.

### rs943080 is associated with regulatory effects on *VEGFA* expression

To examine the potential function of a SNP with a high posterior probability of causality, we further examined the molecular data associated with rs943080 at the *VEGFA* locus. *VEGFA* is a strong candidate for the regulatory target of this SNP as it is highly expressed in RPE and the SNP is within a Hi-C chromatin loop identified in iPSCs (Greenwald et al., 2018) associated with the *VEFGA* promoter (**Figure 4A**). The rs943080 SNP is located ~90KB from the start position of the *VEGFA* gene in a strong iPSC-RPE ATAC-seq peak (**Figure 4B**). While the SNP does not interrupt a transcription factor (TF) motif or appear to create a new motif (data not shown), the peak overlaps three motifs of TFs relevant for the eye and/or RPE (SOX9 (Tang et al., 2015), COUP-TFI/II (Tang et al., 2015), and ZIC3 (Zhang et al., 2004); **Figure 4C**); additionally, all three TFs were expressed in iPSC-RPE (data not shown). In addition, COUP-TFII has been implicated in angiogenesis and regulation of VEGFR in the cardiovascular system (Pereira et al., 1999), suggesting that it could also play a role in angiogenesis in RPE. We first examined whether the ATAC-seq peak showed allele-specific expression at the rs943080 SNP. For the six subjects for which we obtained iPSC-RPE, we used whole genome sequence data to determine that five were heterozygous for the rs943080 SNP (C/T) and one was homozygous for the risk allele (T/T). Across the five heterozygous subjects, we observed a significant allele specific effect (ASE) for the risk allele (P = 3×10^−12^, **Figure 4C**), indicating that the risk allele was associated with stronger chromatin accessibility. These results suggest that rs943080 may act through allele-specific regulation of the distal ATAC-seq peak.

**Figure 4:**
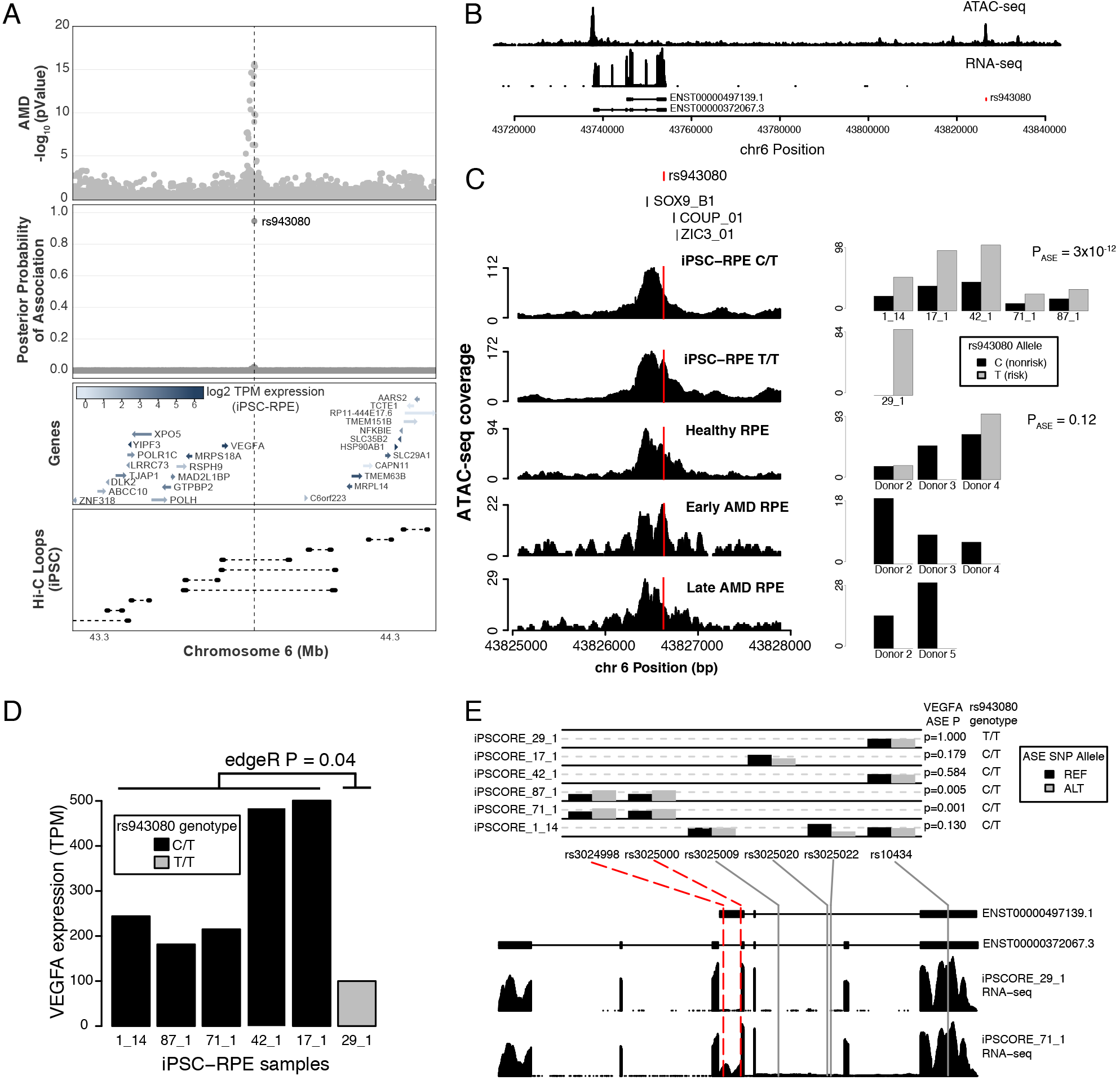
The rs943080 SNP is associated with expression of *VEGFA* and allele-specific expression of a non-coding *VEGFA* transcript. **A)** Genomic region surrounding the genome-wide significant lead SNP near *VEGFA* showing AMD GWAS association (top), posterior probability of causal association from fgwas for rs943080 (middle-top), gene location and expression level in iPSC-RPE (middle-bottom), and Hi-C chromatin loops measured in iPSCs (bottom). **B)** Wig-coverage plots showing example ATAC-seq and RNA-seq data from iPSC-RPE. Two *VEGFA* transcripts are shown and the location of rs943080 is indicated. **C)** rs943080 shows allele-specific expression of an ATAC-seq peak. TF motifs are shown at the top of example wig coverage plots from each type of RPE samples (iPSC-RPE that is heterozygous for rs943080, iPSC-RPE that is homozygous risk for rs943080, RPE from normal adult eyes, RPE from AMD patients with early stage disease, and RPE from AMD patients with late stage disease); the red line indicates the location of rs943080 in the peaks (left). For each sample, the number of reference (non-risk) and alternate (risk) alleles are shown as a barplot; for heterozygous samples, a meta-analyzed P-value of ASE is reported (right). **D)** Barplot showing overall TPM gene expression for *VEGFA* in each of iPSC-RPE samples. Bars are color coded by sample genotype and the edgeR P-value for the difference between genotype groups is shown. **E)** ASE of *VEGFA* in the iPSC-RPE. Barplots showing the allele frequencies for six SNPs covering the *VEGFA* gene in each sample where they are heterozygous; ASE P-values for overall gene expression are listed for each sample on the right (top). The location of each of the SNPs relative to *VEGFA* transcripts and wig coverage plots of RNA-seq data from a homozygote risk (iPSCORE_29_1) and heterozygote (iPSCORE_71_1) individual are shown (bottom); red dotted lines highlight the two SNPs that overlap an exon of a non-coding transcript (ENST00000497139.1) and are driving the significant ASE association in two individuals.

To examine whether adult RPE samples also showed ASE, we inspected allelic depth in the samples from Wang et al. (Wang et al., 2018). We observed reads for both alleles for 2 of the 3 adult healthy RPE subjects, consistent with these individuals being heterozygous for the variant, but did not observe significant ASE in these samples (P = 0.12, potentially, due to low comparative read depth. In the adult early and late-stage AMD RPE samples, we failed to observe any reads with the risk allele, which could be explained because 1) they are all homozygous for the non-risk genotype or 2) in AMD samples, there is strong ASE for the non-risk allele. While we do not have genotype data for these four subjects, given the risk allele frequency (0.52 in European populations, higher in other populations and in AMD subjects), it is unlikely that they would all be non-risk homozygotes (probability = 0.003). Therefore, it is possible that the rs943080 variant could show ASE of chromatin accessibility associated with the nonrisk allele in RPE from AMD subjects, in contrast to the ASE of chromatin accessibility associated with the risk allele in iPSC-RPE and adult healthy RPE. These results therefore suggest that rs943080 is associated with differential chromatin accessibility in iPSC-RPE and healthy adult RPE, but that it may act differently in an AMD context.

To examine whether the rs943080 SNP was also associated with *VEGFA* gene expression, we examined overall expression levels and ASE in RNA from the iPSC-RPE samples. We observed that the five heterozygous samples showed significantly higher overall gene expression than the one homozygous risk sample (edgeR P = 0.04, **Figure 4D**), suggesting that the rs943080 risk allele may drive decreased *VEGFA* gene expression in RPE. We then examined transcriptome data and identified two samples with significant ASE for *VEGFA* RNA (iPSCORE_87_1 and iPSCORE_71_1, **Figure 4E**). To determine whether the ASE was associated with a specific region of the *VEGFA* transcripts, we examined the locations and frequencies of the SNPs that were used to estimate ASE. We found that the ASE for these two subjects was driven by two SNPs that were located in an exon of a non-coding transcript (ENST00000497139.1) and which did not overlap coding exons, suggesting that the ASE observation could be restricted to the non-coding transcript. Consistent with this model, we observed that a third SNP located in the 3’UTR, which is shared across almost all *VEGFA* transcripts, had high coverage in multiple samples and was not associated with ASE. Thus, while rs943080 is associated with an overall decrease in VEGFA expression, it may be mediated not through ASE of all transcripts, but rather through a specific non-coding transcript.

## Discussion

In this manuscript, we have generated RNA-seq, ATAC-seq, and H3K27ac ChIP-seq data from six human iPSC-RPE differentiations as well as data from one human fetal RPE sample. We integrated this data with published ATAC-seq data from RPE and retina from adults with and without AMD to prioritize specific variants at AMD risk loci. The majority of the variants whose prioritizations were altered by our model were located in regulatory regions, although some were also coding variants in genes. Overall, we observed strong improvements in the prioritization of four variants, including a potential regulatory SNP near the *VEGFA* gene.

By examining RNA-seq and ATAC-seq data from individuals with varying genotypes at the rs943080 SNP near *VEGFA*, we suggest that the risk variant may result in decreased gene expression of a non-coding transcript of *VEGFA* in developing or normal RPE cells. While this transcript has not been well characterized, it is highly expressed in many tissues (Gamazon et al., 2018) and could have a regulatory role as it includes the 3’UTR, which has been shown to have regulatory effects on translation (Ray et al., 2009). These results are relevant for the treatment of AMD as anti-VEGF is the only currently approved treatment for “wet” AMD that shows choroidal neovascularization. While this treatment improves symptoms that are driven by the growth of blood vessels, such as swelling and bleeding, it is not clear that it stops progression of the disease, and it may be associated with the further development of geographic atrophy seen in “dry” AMD (Enslow et al., 2016). Specific knockout of *VEGFA* in RPE in mice has been shown to result in dysfunction and loss of the choriocapillaris and cone photoreceptors, suggesting that low levels of *VEGFA* could affect AMD pathology (Kurihara et al., 2012). Our results are consistent with this model and suggest that the rs943080 risk variant could act through a reduction of *VEGFA* expression prior to AMD onset. We were also able to examine the chromatin accessibility of this region in previously published ATAC-seq samples from adult RPE with and without AMD. While we did not have access to genetic data for these subjects, we observed biallelic accessibility in normal RPE samples and monoallelic accessibility in AMD samples with only the non-risk allele expressed. As increased chromatin accessibility in normal cells was associated with decreased *VEGFA* expression, it is plausible that a lack of chromatin accessibility in AMD samples could be associated with
 increased *VEGFA* expression. Further study of the transcriptional effects of this variant in subjects with genotype data are therefore needed.

While we were able to gain insight into the *VEGFA* locus because we had a number of heterozygous individuals as well as an individual that was homozygous for the rs943080 risk allele, additional samples will be required to identify target genes for most GWAS loci. To gain insight into functional genetic variation genome-wide, it will likely be necessary to obtain data for hundreds of samples. As high-quality RPE samples are challenging to obtain from human cadavers and because sample limitations may restrict molecular characterization, iPSC-RPE could potentially be generated from the hundreds of individuals needed. These data suggest that iPSC-RPE are well suited for the genetic characterization of functional variation in RPE and that these large studies could aid in the identification of causal variants at AMD risk loci.

## Experimental Procedures

### RPE derivation and characterization

We obtained iPSC-RPE using a slightly modified version of the Maruotti et al. protocol (Maruotti et al., 2015). The iPSCs were cultured as a monolayer on Matrigel^®^ in mTeSR1 medium. Once cells reached the desired confluency (>80%, approximately 5 days), mTeSR1 medium was replaced with RPE differentiation medium (RPE DM) (24ml/10cm dish) and cells were cultured for 24h. After one day, RPE DM was supplemented with 10mM Nicotinamide (NIC) and 50nM Chetomin (CTM; a strong inducer of RPE (Maruotti et al., 2015)) (RPE DM + NIC + CTM). After 2 weeks, cells were cultured in RPE DM medium supplemented with 10mM NIC (RPE DM + NIC). The cells were then split at day 28 and day 56 with culturing in RPE medium until day 84. About one week after each passage, the polygonal cells formed a very tight, fully pigmented monolayer. Of note, at day 84 we were able to cryopreserve, thaw and then further expand the iPSC-RPE.

### Cellular Data Generation

#### Flow cytometry analysis

RPE were analyzed for ZO-1 and MITF co-expression using flow cytometry. iPSC-RPE cells at day 84 were collected from T150 flasks as described above, filtered using a 70μm strainer and counted. 5×10^6^ cells were fixed and permeabilized using the Fixation/Permeabilization Solution Kit with BD GolgiStop™ (BD Biosciences 554715) following manufacturer recommendations, and resuspended at the concentration of 1×10^7^/ml. 2.5×10^5^ fixed and permeabilized cells were stained in 1X BD Perm/Wash™ Buffer with a Rabbit polyclonal anti-ZO-1 antibody (Abcam, ab59720; 1:10), Mouse monoclonal anti-MiTF antibody (Abcam, ab12039, 1:10), or appropriate class control antibodies for 1h at room temperature followed by a Donkey-anti-Rabbit AlexaFluor 647 conjugated antibody (Abcam, ab150075; 1:200) or Goat-anti-Mouse AlexaFluor 488 conjugated antibody (Thermo Scientific, A-11001; 1:200). Cells were acquired using FACSCanto II (BD Biosciences) and analyzed using FlowJo software V 10.4. The fraction of ZO-1 and MITF positive cells were similar across all six iPSC-RPE lines (mean = 93.3%, range = 85.8-99.4%; and mean = 98.9%, range = 98.0-99.8%, respectively), confirming the robustness of the protocol.

#### Immunofluorescent characterization

Fresh or cryopreserved iPSC-RPE cells at day 84 were thawed and plated on Matrigel^®^ coated glass Millicell EZ SLIDE 8-well glass slides (Millipore, PEZGS0816) and cultured for 10 days in RPE medium. The cells were then fixed with 4% PFA for 10 min at room temperature (RT), washed twice with PBS, 0.1% Tween^®^ 20, incubated for 20min at room temperature in a blocking – permeabilizing solution (PBS, 1% BSA, 0.1% Triton X-100) and stained overnight at 4°C with either a Rabbit polyclonal anti-ZO-1 antibody (Abcam, ab59720; 1:250) and a mouse monoclonal anti-MiTF antibody (Abcam, ab12039, 1:100) or anti-ZO-1 and mouse monoclonal anti-Bestrophin 1 antibody (Novus Biologicals, NB300-164SS; 1:150) or with the appropriate class control antibodies. The next day, cells were washed three times with PBS and subsequently incubated for 1h at RT with a Goat-anti-Mouse AlexaFluor 488 conjugated antibody (Thermo Scientific, A-11001; 1:250) or Donkey-anti-Rabbit AlexaFluor 647 conjugated antibody (Abcam, ab150075; 1:250) and mounted with the ProLong Gold Antifade Reagent with DAPI (Cell Signaling Technologies, 8961). Cells were imaged using an Olympus FlowView1000 confocal microscope and FlowView ASW V03.01.03.03 or V4.2a software at the UCSD Microscopy Core.

### Molecular Data Generation and Processing

#### RNA-seq

Total RNA was isolated using the Quick-RNA Mini Prep Kit (Zymo) from frozen RTL plus pellets, including on-column DNAse treatment step. RNA was eluted in 40 μl RNAse-free water and run on a Bioanalyzer (Agilent) to determine integrity. Concentration was measured by Qubit. Illumina Truseq Stranded mRNA libraries were prepared and sequenced on HiSeq4000, to an average of 40 M 150 bp paired-end reads per sample. RNA-seq reads were aligned using STAR (Dobin et al., 2013) with a splice junction database built from the Gencode v19 gene annotation (Harrow et al., 2012a). Transcript and gene-based expression values were quantified using the RSEM package (1.2.20) (Li and Dewey, 2011) and normalized to transcript per million bp (TPM).

#### ATAC-seq

We performed ATAC-seq using the protocol from Buenrostro et al. (Buenrostro et al., 2013) with small modifications. Frozen nuclear pellets of 5 × 10^4^ cells each were thawed on ice and tagmented in permeabilization buffer containing digitonin. Tagmentation was carried in 25μl using 2.5μl of Tn5 from Nextera DNA Library Preparation Kit (Illumina) for 60 min at 37°C in a thermomixer (500 RPM shaking). To eliminate confounding effects due to index hopping, all libraries within a pool were indexed with unique i7 and i5 barcodes. Libraries for iPSC-RPE were amplified for 12 cycles. Libraries were sequenced to approximately 80M 150bp paired end reads on the HiSeq 4000 (Illumina) platform. ATAC-seq reads were aligned using STAR to hg19. Duplicate reads, reads mapping to blacklisted regions from ENCODE, reads mapping in chromosome other than chr1-chr22, chrX, chrY, and read-pairs with mapping quality Q<30 were filtered. In addition, to restrict the analysis to regions spanning only one nucleosome, we required an insert size no larger than 140 bp, as we observed that this filtering improved sensitivity to call peaks and reduced noise. Peak calling was performed using MACS2 on BAM files with the command ‘macs2 callpeak --nomodel --nolambda --keep-dup all --call-summits -f BAMPE -g hs’, and peaks were filtered by enrichment score (q < 0.01).

### Data analysis

#### RNA-seq analysis

Principal component analysis (PCA) was performed on the 10,000 genes with the most variable expression (i.e. with the highest standard deviation across samples) across 222 iPSC lines, 141 iPSC-CM lines, one human fetal RPE sample and six iPSC-RPE lines. Functional enrichment of genes associated with each PC was performed using goseq (Young et al., 2010) by comparing the 1,000 genes with the highest or lowest loading on the PC to the other 9,000 used for PCA.

#### ATAC-seq analysis

ATAC-seq data was obtained from Wang et al. via GEO (GSE99287) and SRA (SRP107997). Sequencing reads were processed through the same computational pipeline as data generated for this study. The five healthy donors were treated as unique subjects. For the AMD samples, as samples from the same subject could be annotated as early or late depending on the affected status of the eye, the five AMD donors were treated as unique subjects, whether or not the sample was annotated as early or late AMD.

#### TF enrichment

Enrichment for transcription factor motifs was performed with HOMER (Heinz et al., 2010) using HOCOMOCOv11 motifs at P = 0.0001 (http://hocomoco11.autosome.ru/finalbundle/hocomoco11/core/HUMAN/mono/HOCOMOCOv11coreHUMANmonohomerformat0.0001.motif). Sequences flanking the summits identified by MACS2 were examined using findMotifsGenome.pl – size 200 – p 1 – mask. To cluster TF motif profiles, the enrichment P-values were ranked within each sample across all TFs clustered and clustered using hierarchical clustering with the R package pheatmap (Kolde, 2015).

#### Measuring AMD GWAS enrichment within tissue ATAC-seq peaks

To measure the enrichment of AMD GWAS within different tissues, we applied fgwas (Pickrell, 2014) on each set of ATAC-seq peaks independently. Variants were used if their alternate allele frequency could be obtained from the 1000 Genomes Project Phase 3 EUR data by matching the chromosome, position, and annotated alleles. This resulted in a loss of 4% of variants tested. Annotations for exonic, promoter, and UTR regions were obtained from GENCODE (Harrow et al., 2012b) annotations (V19). Annotations for missense and synonymous variants were obtained from the 1000 Genomes Project Phase 3 (Genomes Project et al., 2015) functional annotation. ATAC-seq peak regions or H3K27ac peak regions were called separately for each sample and then merged using bedtools v1.7 (Quinlan and Hall, 2010). To identify variants within each set of ATAC-seq peaks or other annotations for input to fgwas, we used bedtools v1.7 (Quinlan and Hall, 2010). For input to fgwas, we calculated Z scores using the β and its standard error, and ran fgwas using these Z scores and standard errors on consecutive ~1 Mb intervals across the genome (-k 3355).

#### Fine-mapping AMD GWAS loci

To perform fine-mapping of the AMD GWAS loci, we trained an fgwas model containing annotations from ATAC-seq data (Early AMD RPE, Earlt AMD Retina, Late AMD RPE, Late AMD Retina, Healthy RPE, Healthy Retina, human fetal RPE, and iPSC-RPE), H3K27ac data (human fetal RPE and iPSC-RPE), and genome annotations (exons, promoters, untranslated 3’ or 5’ regions, missense variants, or synonymous variants). We applied fgwas on these annotations, and chose the cross-validation penalty to use by first trying penalties between 0.05 and 0.30 in steps of 0.05, and then positive nonzero penalties in 0.01 steps surrounding the best penalty by 0.05, resulting in a tested range of penalties from 0.25-0.30; a final penalty of 0.30 was selected as it had the best likelihood. Finally, we removed annotations from the model until the likelihood stopped increasing, resulting in 11 annotations from the three types being retained: ATAC-seq (Early AMD RPE, Early AMD Retina, Late AMD RPE, Late AMD Retina, Healthy Retina, human fetal RPE, and iPSC-RPE), human fetal H3K27ac, and genome annotations (exons, missense variants, and promoters). We used the model with fgwas to update the Bayes Factors for each variant using the cross-validation estimated ridge parameter and calculated the posterior probability of causality (PP) for each variant within 1 Mb windows flanking the reported lead variant (Fritsche et al., 2016). For two of the lead variants, we were unable to obtain allele frequencies from the 1000 Genomes Project, resulting in them being removed from the analysis (TRPM3/rs71507014 and MMP9/rs142450006). We therefore did not perform fine-mapping for these loci. The PPA is the proportion of the total GWAS risk signal at a locus measured by Bayes Factors that is attributed to a particular variant, multiplied by the probability that the genomic region contained a real signal (set to 1 for GWAS loci). Variants that were associated with annotation information were classified into functional classes based on the following criteria in the following order: Coding = missense variant, Local Regulatory = promoter annotation at least one ATAC-seq or H3K27ac annotation, Distal Regulatory = at least one ATAC-seq or H3K27ac ChIP-seq annotation, Unknown = all others.

#### *VEGFA* locus annotation

To visualize the *VEGFA* region (Figure 4A), −log10 P-values from (Fritsche et al., 2016) were plotted along with the PPA of all SNPs after prioritization with fgwas. TPM normalized RNA-seq expression data from an iPSC-RPE (iPSCORE_1_14) was used to identify expressed genes in the region. Hi-C loops were obtained from Greenwald et al. (Greenwald et al., 2018).

#### ATAC-seq peak ASE of rs943080

To examine ASE at the ATAC-seq peak containing rs943080, allelic read depth was measured using samtools (Li, 2011) mpileup. Read depth from all samples associated with the same subject were combined. For the iPSC-RPE, the genotype at rs943080 was obtained from whole genome sequence data (dbGaP phs001325). For each subject, an ASE P-value was calculated by testing the read depth counts to the expected frequency (50%) using a binomial test (binom.test in R). Across all heterozygotes, a meta-analysis P was calculated using the “sumlog” method in the R package metap (Dewey, 2018). For adult healthy subjects, only the donors with at least 2 reads from each allele were considered heterozygotes and the meta-analysis P was calculated in the same manner as the iPSC-RPE. For the adult early and late-stage AMD samples, since we did not observe any samples with alternate reads, we calculated the probability of observing four subjects with a homozygous reference genotype as the square of the reference allele frequency in the 1000 Genomes Project (Genomes Project et al., 2015) EUR population (0.48) to the fourth power.

#### RNA-seq analysis at *VEGFA*

Expression differences between the iPSC-RPE from the five rs943080 heterozygous subjects were compared to the one homozygous subject using edgeR (Robinson et al., 2010). Reads counts were merged across all transcripts to examine gene-level differences. The unadjusted quasi-likelihood F-test P-value is reported, although the likelihood ratio test result was similar. ASE analysis of RNA-Seq data was performed using MBASED (Mayba et al., 2014). Heterozygous SNVs were identified by intersecting variant calls from WGS with exonic regions from Gencode v19. The WASP pipeline (van de Geijn et al., 2015) was employed to reduce reference allele bias at heterozygous sites. The number of read pairs supporting each allele was counted using the ASEReadCounter from GATK (3.4-46) (DePristo et al., 2011). Heterozygous SNVs were then filtered to keep SNVs where the reference or alternate allele had more than 8 supporting read pairs, the reference allele frequency was between 2-98%, and the SNV was located in unique mappability regions according to wgEncodeCrgMapabilityAlign100mer track, and not located within 10 bp of another variant in a particular subject (heterozygous or homozygous alternative) (Consortium, 2015; Lappalainen et al., 2013; Mayba et al., 2014). The raw gene-based P-values were reported. For visualization, only SNPs with a combined read depth of more than 30 were shown.

## Author Contributions

Conceptualization, E.N.S., A.D.C., and K.A.F.

Software, E.N.S., M.D., W.W.G., and H.M.

Formal Analysis, E.N.S., M.D., and W.W.G.

Investigation, A.D.C., V.B., L.R.A., and S.B.

Data Curation, E.N.S., M.D., and H.M.

Writing – Original Draft, E.N.S, A.D.C., M.D., W.W.G., and K.A.F.

Writing – Review & Editing R.A., S.B., and R.P.

Visualization, E.N.S., M.D., and W.W.G.

Supervision, E.N.S., R.P., A.D.C., S.B., and K.A.F.

Project Administration, E.N.S., A.D.C., and K.A.F.

Funding Acquisition, S.B., and K.A.F.

## Acknowledgments

This work was supported in part by a California Institute for Regenerative Medicine (CIRM) grant GC1R-06673 and NIH grants HG008118, HL107442, DK105541, DK112155 and EY021237. RNA-seq were performed at the UCSD IGM Genomics Center with support from NIH grant P30CA023100. W.W.G. was supported by the National Heart, Lung, And Blood Institute of the National Institutes of Health under Award Number F31HL142151. L.R.A. was funded by a scholarship from the Brazilian Coordenação de Aperfeiçoamento de Pessoal de Nível Superior (CAPES). S.B. was funded by a Fulbright-Fight for sight scholarship and a Foundation Fighting Blindness Career Development Award.

## Accession Numbers

ATAC-seq and RNA-seq data can be accessed through dbGaP (BAM files, phs000924), the Gene Expression Omnibus *(in progress)*, and through the UCSC Genome Browser (iPSCORE_RPE Public Track Hub).

## Supplemental Table Descriptions

Table S1: List of samples and data identifiers used in the study. Related to Figure 1.

Table S2: GO term results for analyses of PCs 3 and 4. Related to Figure 2.

Table S3: Full list of – ln P-values for transcription factor motif enrichment. Related to Figure 2.

Table S4: Full list of lead variants prioritized by fgwas. Related to Figure 3.

